# Real-time, automated, standardized, and transparent analysis of microfluidic nanoparticle data with RPSPASS

**DOI:** 10.64898/2026.03.30.715405

**Authors:** Michelle L. Pleet, Sean Cook, Bryce Killingsworth, Tim Traynor, Dove-Anna Johnson, Emily H. Stack, Verity Ford, Cláudio Pinheiro, Jessie E Arce, Jason Savage, Matthew Roth, Aleksandar Milosavljevic, Ionita Ghiran, An Hendrix, Steven Jacobson, Joshua A. Welsh, Jennifer C. Jones

## Abstract

Extracellular vesicles (EVs) are lipid spheres released from cells. Research utilizing EVs has met several hurdles owing to the small size of the majority of EVs and other nanoparticles (<150 nm) and the lack of detection technologies capable of providing high-throughput single particle measurements at this scale. The use of high-throughput single particle measurements is critical for the assessment of EV heterogeneity and abundance which are features often used to assess the development of isolation protocols or particle characterization. The Coulter principle, known in the field as resistive pulse sensing (RPS), has been used for several decades to size and count cells. More recently, this technology has evolved to accommodate nanoparticle analysis. In the last decade a platform utilizing microfluidic resistive pulse sensing (MRPS) has been demonstrated for nanoparticles, offering ergonomic characterization of nanoparticles along with utilizing open format data. To date, assessment of MRPS accuracy and reporting standards have not been assessed. With the aim of increasing data accuracy, ergonomics, and reporting transparency, we developed a microfluidic resistive pulse sensing post-acquisition analysis software (RPSPASS) application for automated cohort calibration, population gating, statistical output, QC plot generation, alternative data file outputs, and standardized reporting templates.

## INTRODUCTION

Extracellular vesicles (EVs) are the focus of an increasing area of research with respect to developing diagnostic, prognostic, and therapeutic biomarkers along with understanding basic science.^1^ Characterizing small EVs (<150 nm), however, remains a hurdle resulting in the need for the development of novel and more sensitive detection technologies.^2^ Due to the lack of gold-standard detection technologies for simultaneous size and concentration measurement, the use of orthogonal methods with non-overlapping technologies (e.g. optical *vs*. non-optical) has been recommended, with each method ideally being able to define their sensitivity in traceable standard units.^3^

A non-optical method proving beneficial for characterizing EV size and concentration at a single particle level is microfluidic resistive pulse sensing (MRPS), commercially available as the nCS1 platform, Spectradyne.^4, 5^ MRPS is based on the Coulter principle, whereby particle sizing and counting is based on measurable changes in electrical impedance produced by nonconductive particles suspended in an electrolyte.^6^ This methodology was popularized in the EV field in the form of tunable resistive pulse sensing (TRPS) with the commercial qNano platform, Izon Science.^7–9^ MRPS offers advantages over TRPS in its implementation by having a design that is more resistant to clogging from larger particles and only requiring loading of a sample onto a microfluidic chip that can be discarded post-acquisition. Furthermore, the implementation of open-source files makes it possible to inspect and apply further data analyses.

MRPS offers utility as an orthogonal metric to optical methods such as flow cytometry and nanoparticle tracking methods. Its ability to provide single particle diameter and concentration measurements down to the ∼55 nm diameter range are of great utility for EV and nanoparticle characterization. Furthermore, traceable size standards can be used to calibrate and define the limit of sensitivity for RPS methodologies, making them one of the most reliable and reproducible characterization methods for high-throughput single particle quantification.

A current limitation to analyzing and reporting MRPS data is automating the removal of acquisition periods where transient clogging has occurred, dynamically calibrating diameter axes where necessary, and exporting files in a format that is compatible with third-party gating software and online repositories. Here, we demonstrate the use of resistive pulse sensing post-acquisition analysis software (RPS_PASS_). RPS_PASS_ is a free, open-source software application that dynamically calibrates diameter and concentration measurements, removes outliers across sample acquisition, performs automated sample and cohort gating, spike-in bead removal, and exports files in numerous file formats. One of these file formats is the .fcs file format that enables the use of gating tools in well-established flow cytometry software, along with compatibility in sharing to online repositories for reporting (i.e., FlowRepository and exRNA atlas).^10–13^

## MATERIALS AND METHODS

### Protein measurements

BSA concentration was measured using a NanoDrop 2000 Spectrometer (Thermo Fisher Scientific, USA). Prior to recording concentration using the NanoDrop, the sensor was rinsed with deionized water and dried with a cotton bud before a baseline reading was taken using DPBS. Two microliters of sample were then placed on the sensor and a concentration reading was recorded three times. Recordings were exported to .xlsx files. Recordings with negative readings were assumed to be 0 µg mL^-1^.

### Cell culture

The immature dendritic cell line DC2.4 was kindly provided by Kenneth Rock (University of Massachusetts Medical School, Boston, MA) and cultured in phenol red-free RPMI-1640 medium supplemented with 10% EV-depleted FBS, 1% L-glutamine, 1% penicillin-streptomycin and 0.1% β-mercaptoethanol (ThermoFisher). For EV-depleted FBS preparation, 20% FBS containing RPMI was ultracentrifuged for 18 hours at 100,000 x *g* at 4°C in a 45Ti fixed angle rotor using polycarbonate tubes (both from Beckman Coulter). After ultracentrifugation, the top 50 mL of medium suspension was harvested, filtered with 0.2 µm PES filter bottles, and stored at 4°C. To produce DC2.4-derived EVs, cells were cultured for 2–3 days in EV-depleted media with supernatants harvested before confluence was reached. EV-containing cell culture supernatants were aspirated from tissue culture flasks, transferred to 50 mL tubes, and centrifuged at 2000 x *g* twice for 10 minutes. Cell-free supernatants were aliquoted into 60 mL JumboSep canisters (PALL Corporation) and concentrated using 100 kDa filters (JumboSep, PALL Corporation) until ∼5 mL remained. The 5 mL of cell-free concentrates were then run on a 10 mL 70 nm size exclusion column (qEV10, Izon Science) with 5 mL fractions collected. The first 15 fractions were collected starting immediately after the sample was loaded onto the column. EV concentration and protein content of fractions were approximated using nanoparticle tracking analysis (NanoSight LM10, Malvern) and NanoDrop (Thermo Fisher Scientific), respectively (**Fig. 1A**). Fraction 7 was used for downstream characterization. TEM of isolated EVs was performed to confirm purity of preparations and as a method of EV diameter characterization for comparison (**Fig. 1B**).

**Figure 1:**
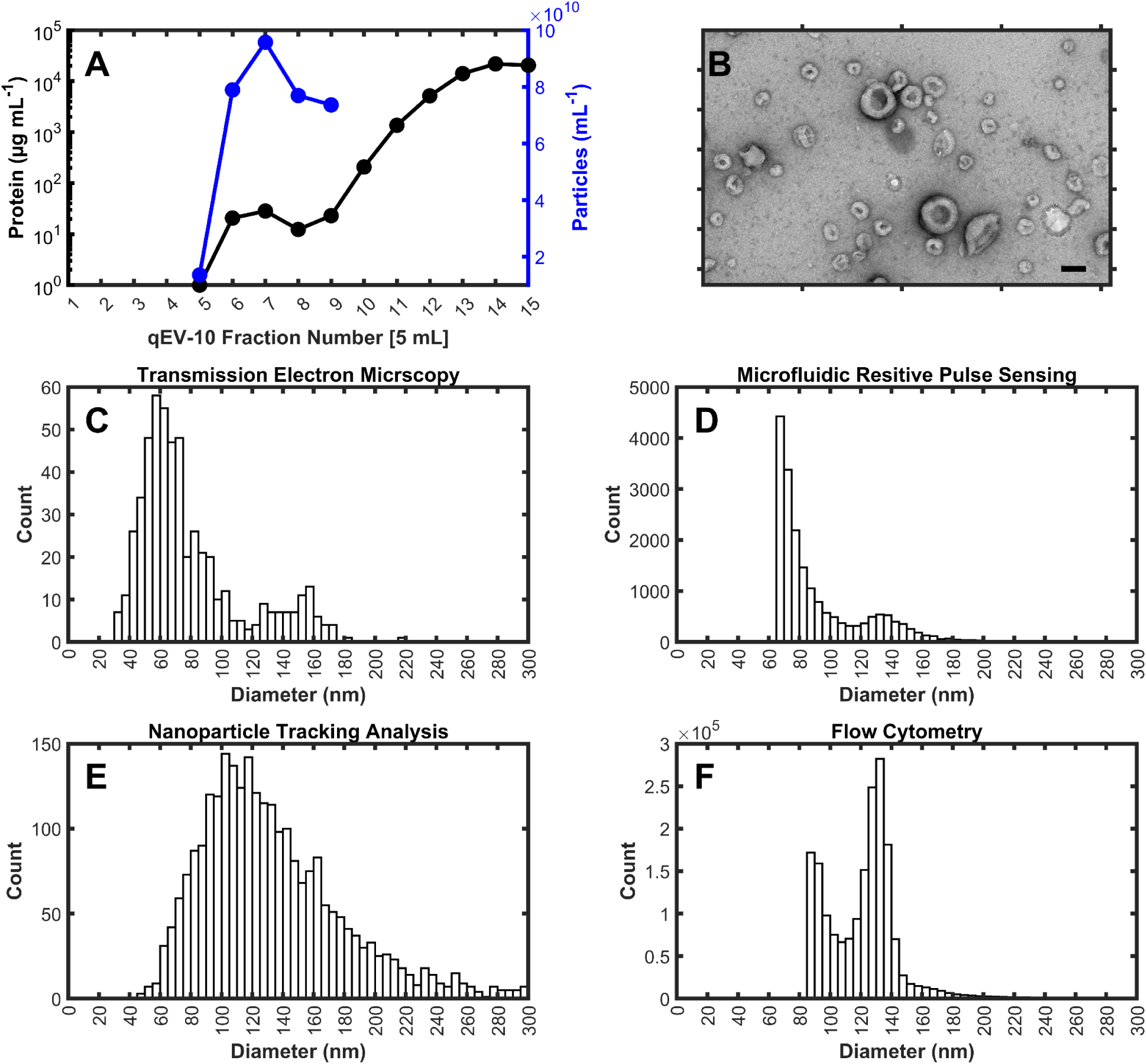
Cross-platform characterization of DC2.4 EV diameter distribution. (**A**) Measurement of size exclusion chromatography fraction protein concentration (black line) and number concentration (blue line). (**B**) Representative transmission electron microscopy image of negatively stained DC2.4 EVs from Fraction 7. Scale bar represents 100 nm. (**C**) Diameter distribution of fraction 7 DC2.4 EVs using transmission electron microscopy and negative staining. (**D**) Diameter distribution of fraction 7 DC2.4 EVs using microfluidic resistive pulse sensing. (**E**) Diameter distribution of fraction 7 DC2.4 EVs using nanoparticle tracking analysis. (**F**) Diameter distribution of fraction 7 DC2.4 EVs using flow cytometry and light scatter calibration assuming average refractive index (wavelength; 405 nm, core RI; 1.38, membrane RI; 1.48, membrane thickness; 6 nm. All diameter distributions were binned with 5 nm increments.

### Cerebrospinal fluid (CSF) isolation and storage

CSF samples were obtained by lumbar puncture in the clinical center at NIH. After centrifugation at 1300 x *g* for 10 minutes, the supernatants were collected into cryotubes and immediately frozen at -80 ℃ until use. Samples for MRPS were subjected to only a single freeze-thaw before use.

### Ethics statement

CSF samples used in this study were collected from the subjects followed at the National Institute of Neurologic Disorders and Stroke under protocols # 98-N-0047, 89-N-0045, 13-N-0017, 13-N-0149. Prior to study inclusion, written informed consent was obtained from the subjects in accordance with the Declaration of Helsinki.

### Flow cytometry of EVs

EV analysis was carried out using a Cytek Aurora (Cytek Biosciences), configured with 4 lasers (405, 488, 561, 640 nm) and an enhanced small particle (ESP) 405 nm detector. Diameter was calculated for EVs using FCM_PASS_ software (v3.03).^14^ Light scatter parameters were calibrated into units of diameter by assuming a high EV refractive index. Full calibration details can be found in the MIFlowCyt-EV report, **Electronic Supplementary Information 1**.^15^ Quality control plots of rEV detection can be found in **Supplementary Figures 1 & 2**.

### Nanoparticle tracking analysis (NTA)

Detectable sample concentration was approximated using a NanoSight LM10 instrument (Malvern, UK), equipped with a 405 nm LM12 module and EMCCD camera (DL-658-OEM-630, Andor). Video acquisition was performed with NTA software v3.4, using a camera level of 14. Three 30 second videos were captured per sample. Post-acquisition video analysis used the following settings: minimum track length = 5, detection threshold = 4, automatic blur size = 2-pass, maximum jump size = 12.0. Exported datasets were compiled and plotted using scripts written in MATLAB (v9.7.0.1261785 (R2019b) Update 3, The Mathworks Inc, Natick, MA).

### MRPS acquisition

MRPS analysis of samples was carried out using the Spectradyne nCS1™ (Spectradyne LLC, USA). Our study focused on the use of TS-400 microfluidic cartridges which have an advertised measurement range of 65 to 400 nm.

### Recombinant EV generation and isolation

Generation and isolation of recombinant EVs (rEVs) was performed as previously described.^16, 17^ In brief, HEK293T cells were seeded in Falcon cell culture Multi-Flasks (cat. no. 353144, Corning, USA) and transiently transfected with pMET7-gag-EGFP plasmid at 70-80% confluency using 25 kDa linear polyethyleneimine (PEI) (cat. no. 23966, Polysciences, USA). Forty-eight hours following transfection, cells were washed three times using Opti-MEM (cat. no. 31985054, ThermoFisher) followed by a 24 hour incubation in Opti-MEM supplemented with 100 IU mL^-1^ penicillin and 100 mg mL^-1^ streptomycin at 37°C and 10% CO2. The conditioned medium (CM) was collected and centrifuged for 10 min at 300 x *g* and 4°C, passed through a 0.45 µm cellulose acetate filter (cat. no. A35999, NovoLab, Belgium), and concentrated to 1 mL using a Centricon Plus-70 centrifugal filter device (cat. no. UFC701008, Merck, Germany). Next, rEVs were isolated from the concentrated CM (CCM) by OptiPrep velocity gradient (OVG). The velocity gradient was prepared by layering 11 iodixanol solutions (cat. no. AXI-1114542, Axis-Shield, Norway) of 1.3 mL, ranging from 18 to 6% with 1.2% decrements, from the bottom to the top in a 16.8 mL open top polyallomer tube (cat. no. 337986, Beckman Coulter, USA) using the Biomek 4000 automated workstation (Beckman Coulter). One milliliter of CCM was overlaid on top of the gradient that was then centrifuged for 1 hour and 56 min at 186,700 x *g* and 4°C (SW 32.1 Ti rotor, Beckman Coulter). Gradient fractions of 1 mL were collected from top to bottom using the Biomek 4000 automated workstation. Fractions 10, 11, 12 and 13 corresponding to buoyant density of 1.076-1.088 g mL^-1^, were collected, pooled, and diluted to 16 mL in PBS and centrifuged for 3 hours at 100,000 x *g* and 4°C (SW 32.1 Ti rotor, Beckman Coulter). The resulting pellet was resuspended in PBS containing 5% trehalose (cat. no. T0167, Merck, Germany) and stored at 4°C until further use.

### Sample preparation for MRPS

Commercial and laboratory-generated rEVs (Sigma, Cat. SAE0193) were used for the development of RPS_PASS_. Samples were diluted to a concentration of ∼5×10^9^ particle mL^-1^ in PBS + 1% Tween 20 with 240 nm NIST-traceable beads (3000 series, Thermo Fisher Scientific) at a concentration between 5×10^9^ - 5×10^7^ particle mL^-1^, specified in relevant figures. CSF samples were diluted at 9 parts to 1 only with 240 nm NIST-traceable beads suspended in 0.1 µm filtered DPBS + 1% Tween 20 at a final concentration of 1×10^9^ particles mL^-1^.

### Software development

RPS_PASS_ was developed using MATLAB (v 9.8.0.1323502 (R2022a, Mathworks Inc) with the code available at: https://github.com/CBIIT//RPSPASS.

### Statistical analysis & data availability

CSF particle concentration group comparisons were performed using a Wilcoxon signed-rank test. All data and analysis scripts can be accessed at the following link: https://figshare.com/s/bcd06197af644df99430.

## RESULTS

### Utility of MRPS measurements

To investigate the utility of MRPS measurements for the characterization of EVs as compared to other methods, dendritic cell line (DC2.4)-derived EVs were characterized using transmission electron microscopy (TEM), MRPS, nanoparticle tracking analysis (NTA) and flow cytometry (FCM) (**Fig. 1C-F**). Only TEM was able to fully resolve the bi-modal DC2.4 EV distribution, with the majority of EVs appearing between 40-100 nm in diameter. MRPS detected particles down to 65 nm, with the diameter distribution being concordant with TEM diameter distributions. FCM was limited to a sensitivity of 85 nm, with the larger of the bimodal distribution fully resolved and the diameter distribution obtained using light scatter calibration consistent with TEM and RPS. The diameter distribution from NTA was inconsistent with the three other methodologies, and no clear limit of detection could be defined due to the multitude of parameters required to obtain diameter from NTA.

### Precision and trueness of measurements

While microfluidic cartridges are pre-calibrated in batches, a variety of variables in microfluidic chip production can result in chip-to-chip variation. As an example, NIST-traceable beads (81, 152, 203 nm) were analyzed and overlayed with their certified diameter distributions (**Fig. 2A**). Raw outputs from pre-calibrated MRPS chips were close to certified values, however, they were improved with dynamic diameter calibration (**Fig. 2B**). It was also found that increasing the precision of calculation of the MRPS diameter conversion from arbitrary units, the linearization of diameter data from small to large diameters was improved (**Fig. 2C**). While the results had a high degree of precision, the trueness of their measurements were noticeably inaccurate prior to further calibration (**Table 1**). Measurement drift in some instances may also be observed as a gradual change in diameter distribution of a population over time (**Fig. 2D**). These are likely due to changes in differential pressure that can be introduced from sample or instrumental factors. While applying a single calibration factor to data can result in correcting the trueness of the median diameter measurement, it is not able to correct for measurement drift resulting in increased sample measurement error.

**Figure 2:**
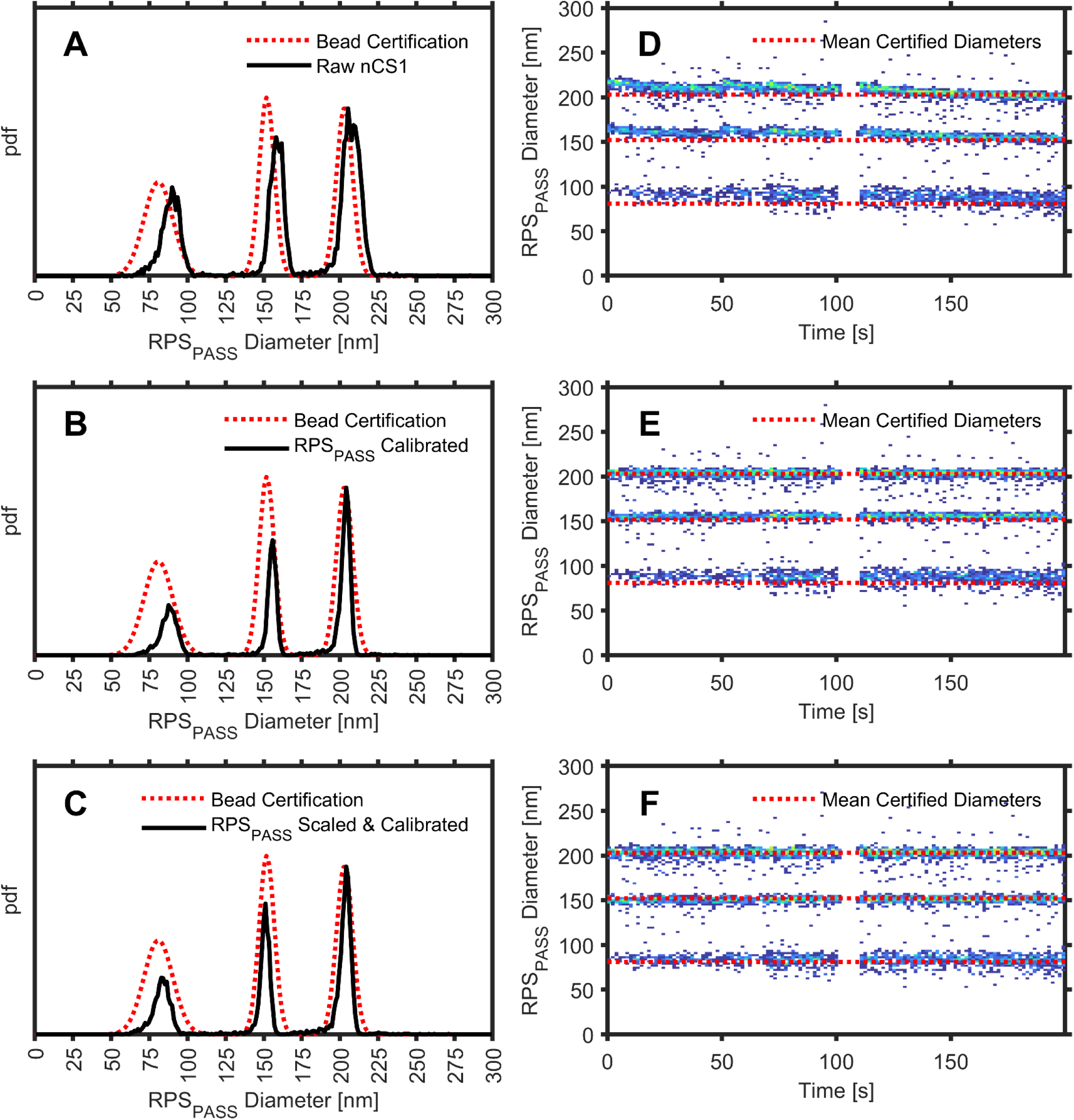
Determining measurement reliability. **(A**) Default nCS1 data for 81, 152, 203 nm NIST traceable beads at 1×10^9^ (black) are overlaid with bead certification specifications (red). Median bead diameters are overlaid (red). (**B**) Dynamically calibrated RPS_PASS_ data using default nCS1 conversion precision for 81, 152, 203 nm NIST traceable beads at 1×10^9^ (black) are overlaid with bead certification specifications (red). Median bead diameters are overlaid (red). (**C**) Dynamically calibrated RPS_PASS_ data using RPS_PASS_ high precision diameter conversion for 81, 152, 203 nm NIST traceable beads at 1×10^9^ (black) are overlaid with bead certification specifications (red). Median bead diameters are overlaid (red). (**D**) Default nCS1 data for 81, 152, 203 nm NIST traceable beads at 1×10^9^ versus acquisition time. (**E**) Dynamically calibrated RPS_PASS_ data using default nCS1 conversion precision for data for 81, 152, 203 nm NIST traceable beads at 1×10^9^ versus acquisition time. (**F**) Dynamically calibrated RPS_PASS_ data using RPS_PASS_ high precision diameter conversion for 81, 152, 203 nm NIST traceable beads at 1×10^9^ versus acquisition time.

**Table 1.** Comparison of NIST-traceable data from certificates, default nCS1 and RPS_PASS_ processing.

| Certified Diameter (nm) | nCS1 Scaling,<br>No Calibration | nCS1 Scaling,<br>RPS<sub>PASS</sub> Calibration | RPS<sub>PASS</sub> Scaling,<br>RPS<sub>PASS</sub> Calibration |
| --- | --- | --- | --- |
| 81±9.5 | 88.5±6.8 | 87.1±6.5 | 82.7±6.3 |
| 152±5.0 | 158.1±5.5 | 155.1±4.5 | 150.7.1±4.8 |
| 203±5.3 | 206±6.5 | 202.9±4.9 | 202.8±5.1 |

To reduce measurement error due to chip-to-chip calibration variation and measurement drift, we developed an automated dynamic calibration algorithm to identify a spike-in bead within a sample and calibrate in acquisition time intervals; by default, this is set to the instrument acquisition interval (default of 10 seconds) (**Fig. 2E-F**). The use of dynamic calibration within RPS_PASS_ resulted in median values having less than 3 nm of difference between measured and certified bead values (**Table 1**). Not only were mean values (82.7, 150.7, 202.8 nm) closer to certified NIST-traceable bead values (81, 152, 203 nm), but their standard deviation was also decreased by being able to account for measurement drift (**Table 1**).

### Optimizing spike-in concentration for software analysis

The use of dynamic calibration requires the use of a spike-in bead while a sample is being run. To avoid uncertainty between sample and bead events, there should be little overlap between the populations. However, the spike-in bead population must be small enough that it will not result in the clogging of a pore. This optimal spike-in diameter range is typically 60-70% of the advertised maximum diameter, and in the case of the TS-400 chips we used beads in the range of 240-280 nm. To demonstrate this, commercially available rEVs were measured at a concentration of 1×10^9^ particles mL^-1^ with 240 nm NIST-traceable spike-in beads at 1×10^9^, 5×10^8^, and 1×10^8^ particles mL^-1^ (**Fig. 3A-F**). The RPS_PASS_ software was successfully able to differentiate the spike-in beads from other events when the spike-in concentration was above 2.5×10^8^ particles mL^-1^. However, at a concentration below 2.5×10^8^ the software was unable to reliably differentiate beads from background events within the 10 second interval window. At concentrations below this, a larger interval window can be utilized, or a single calibration factor can successfully be applied in lieu of dynamic calibration. Using a single calibration factor the software was able to recognize a spike-in bead population and normalize events at a concentration of 1.25×10^8^ particles mL^-1^ (**Fig. 3G-H**), however, this is not recommended.

**Figure 3:**
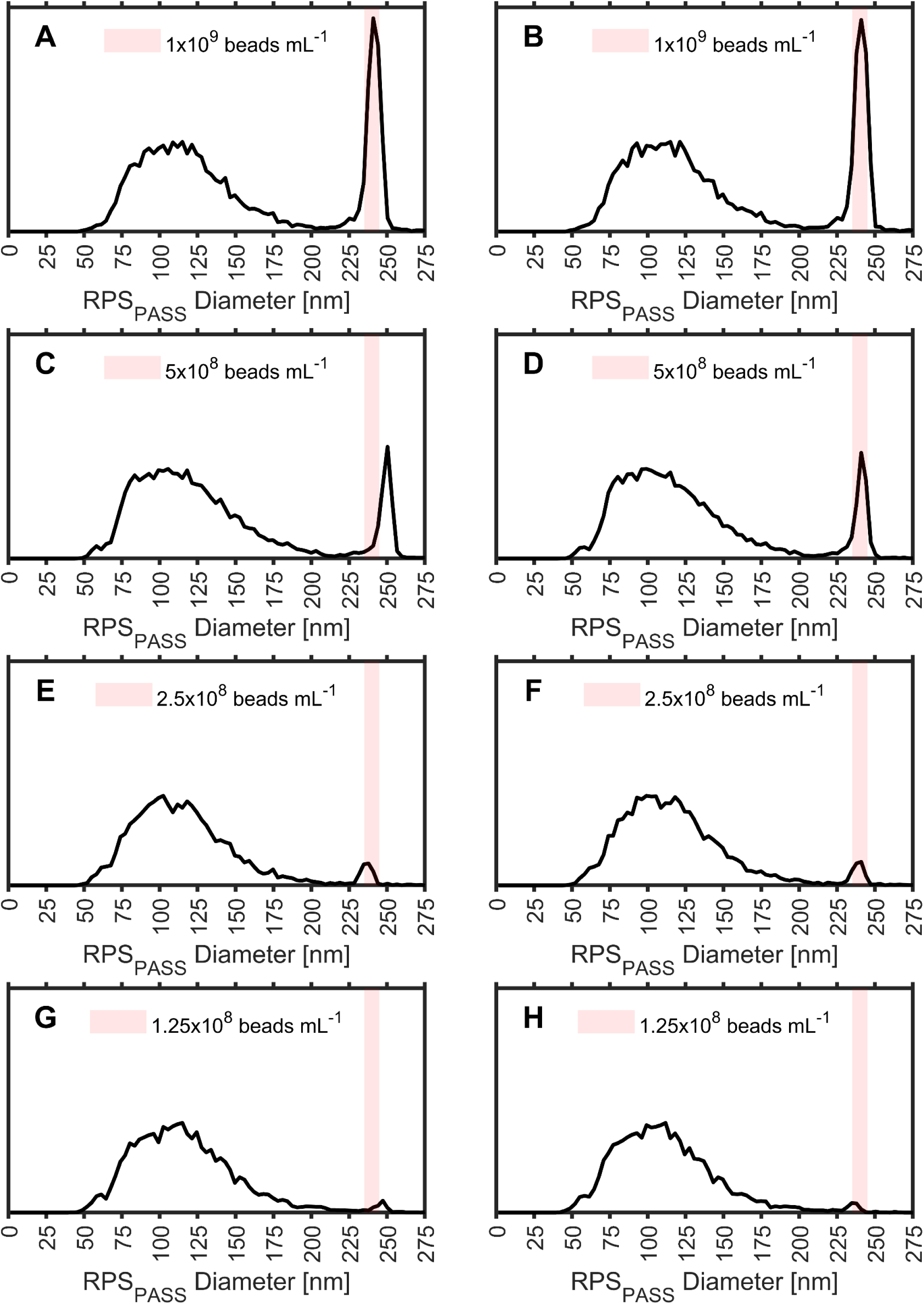
Identifying optimal spike-in concentration for RPS_PASS_. Default nCS1 data for rEVs, with NIST traceable beads spiked-in at various concentrations (**A**, **C**, **E**, & **G**). Dynamically calibrated RPS_PASS_ data for rEVs, with NIST traceable beads spiked-in at various concentrations (**B** & **D**). Non-dynamic calibrated RPS_PASS_ data for rEVs, with NIST traceable beads spiked-in at 5×10^7^ particles mL^-1^ (**F** & **G**). NIST-traceable spike-in bead (235-245 nm) range overlaid in all plots (red).

Chip-to-chip variation, amongst other factors, can result in flow rate variation. Effects such as transient clogging can also impact variation within sample acquisition. Using a batch of acquired data from TS-400 chips the total acquisition volumes per 10 second interval were compared (**Fig. 4A**). Our results showed that most acquisition volumes fell between 10 and 40 pL per 10 second acquisition. While this 4-fold difference in acquisition volume is small and can be accounted for, it does have an impact on chosen spike-in and sample concentrations to allow efficient event acquisition and the ability of the RPS_PASS_ software to recognize the spike-in bead events (**Fig. 4B**). At the median volume of 26 pL per 10 second acquisition, 1×10^8^ detectable particles mL^-1^ will result in 2 recorded events. This increases to 26 events with 1×10^9^ detectable particles mL^-1^ (**Fig. 4B**). To account for chip-to-chip fluidic variation and the ability of RPS_PASS_ software to recognize at least 10 spike-in bead events per 10 second acquisition, we recommend using a spike-in concentration of >8×10^8^ particles mL^-1^ (**Fig. 4B**). Using a single calibration factor with a minimum target of 50 events per sample acquisition allows 5×10^7^ particles mL^-1^ to be sufficient, provided ∼400 seconds of total acquisition (**Fig. 4C**). When the target threshold for individual acquisitions is not met, a single calibration factor is defaulted to by RPS_PASS_ in the analysis. However, applying a single calibration factor is not capable of reducing variation due to measurement drift between acquisitions. Therefore, use of a single calibration factor will improve the trueness of measurement but not increase the precision.

**Figure 4:**
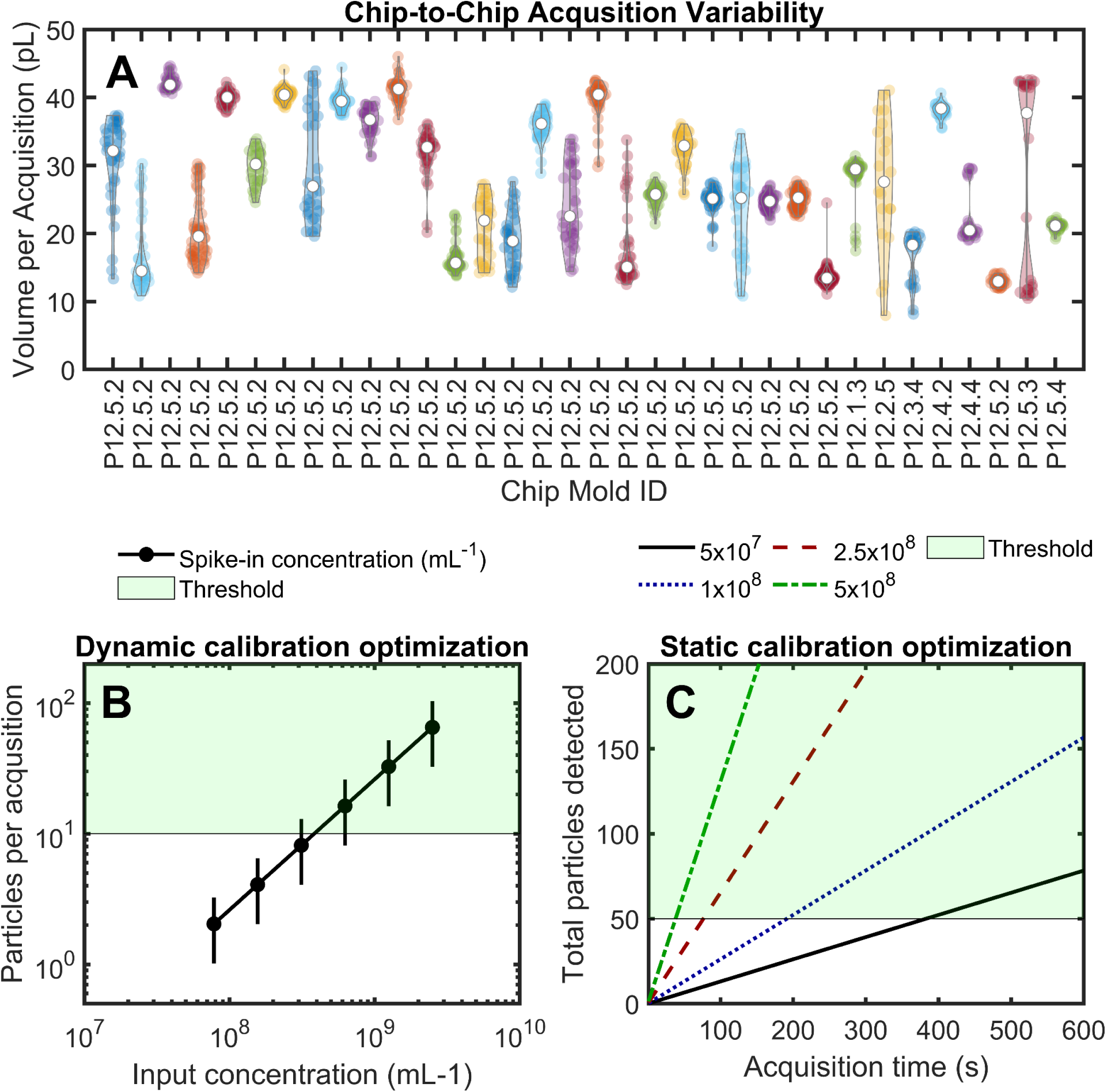
Chip-to-chip flow rate variation. (**A**) Representative data from MRPS mold ID versus total acquisition volume in 10 seconds. (**B**) Particles recorded per 10 second acquisition given a set input concentration assuming the median flow rate from plot A. The RPS_PASS_ threshold for spike-in events per acquisition to achieve dynamic calibration is shown in green. (**C**) Particles recorded per total acquisition time given a set input concentration assuming the median flow rate from panel A. The RPS_PASS_ threshold for spike-in events per acquisition to achieve static calibration is shown in green.

### Development of automated analysis pipeline

Gating particles of interest from background noise and spike-in events can be a labor-intensive task to perform manually over a cohort. Gating particles of interest in a standardized fashion to ensure a fair comparison between samples may also be difficult to achieve, depending upon the software tools available. Over the course of a sample run using MRPS, transient clogs or changes in differential pressure may result in inconsistencies that make gating a cohort even more difficult. We therefore developed an automated outlier detection and gating pipeline within RPS_PASS_ (**Fig. 5-6**). This analysis pipeline incrementally tests every combination of gating parameters to find a set where the recording is most consistent. These outlier dynamic gating parameters include separation index (**Fig. 5A**), spike-in %CV (**Fig. 5B**), particle number to spike-in number (**Fig. 5C**), spike-in transit time (**Fig. 5D**), signal-to-noise transit time ratio (**Fig. 5E**), and sample injection pressure (**Fig. 5F**). Background noise is removed from data using the signal-to-noise transit time ratio > 1 (**Fig. 5G-I**). This signal-to-noise transit time ratio is use solely for individual sample gating in order to maximize detected events (**Fig. 5H**). Alternatively, a standardized gating strategy that increases chip-to-chip comparison accuracy, and thereby allows for cohort gating, utilizes a diameter gate above the noise defined by the signal-to-noise transit time ratio <1 (**Fig. 5I**). The result of gated data from all events (**Fig. 5J**), remaining noise (**Fig. 5K**), and non-noise events (**Fig. 5L**) can then be observed.

**Figure 5:**
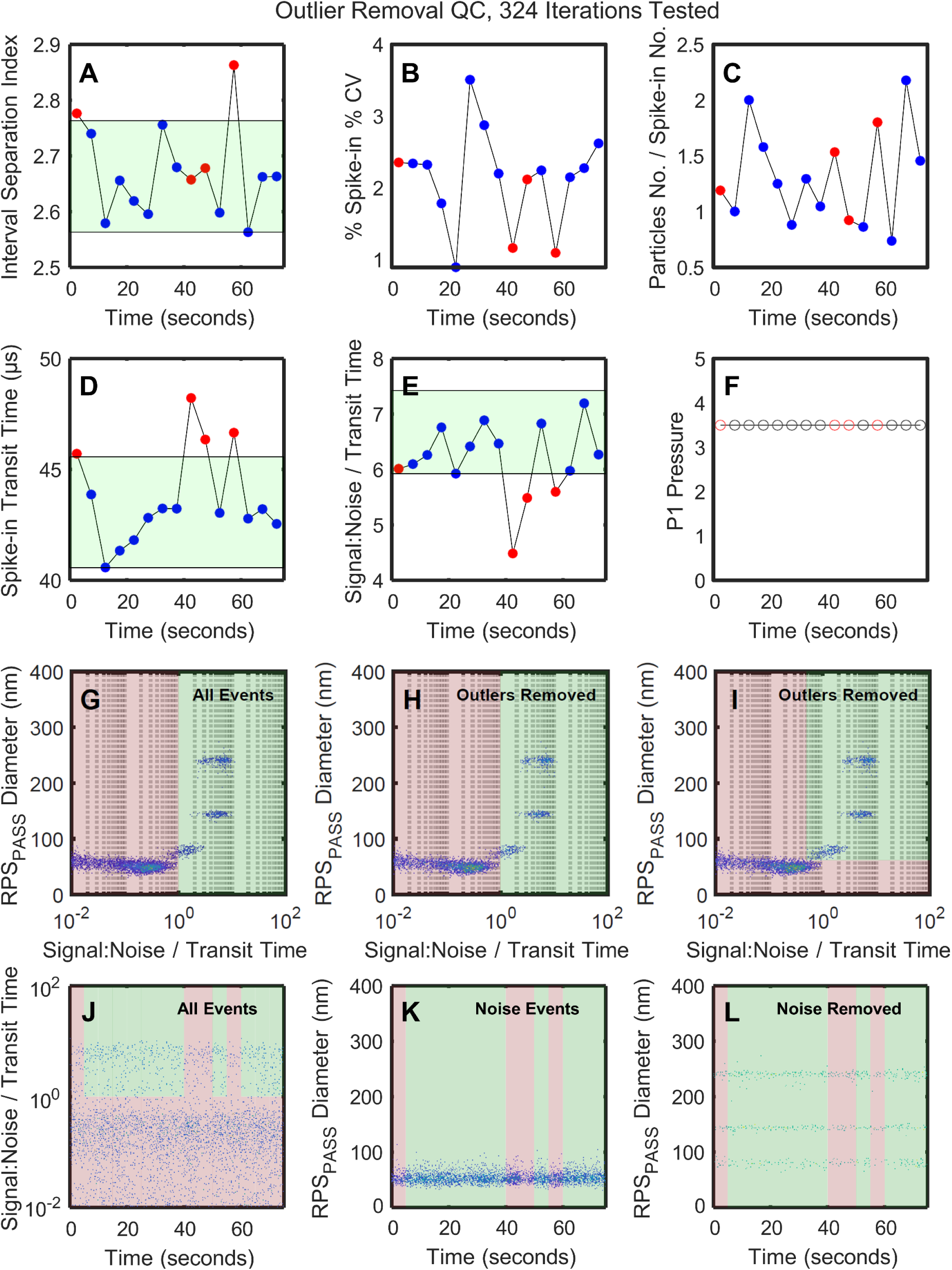
RPS_PASS_ outlier removal debug plot. The summary statistics for each 5 second acquisition are shown vs separation index (**A**), spike-in %CV (**B**), particle number / spike-in number (**C**), spike-in transit time (**D**), signal-to-noise / transit time (**E**), and P1 sample pressure (**F**). The gating of non-background events (green) from background events (red) using the signal-to-noise / transit time parameter with all events (**G**), after outlier removed event (**H**), and with outlier events removed with a standard gating strategy (**I**). The resulting identification of outliers (red) from kept acquisitions (green) are shown (**J-K**), with all events (**J**), noise only events (**K**), and non-noise events (**L**).

**Figure 6:**
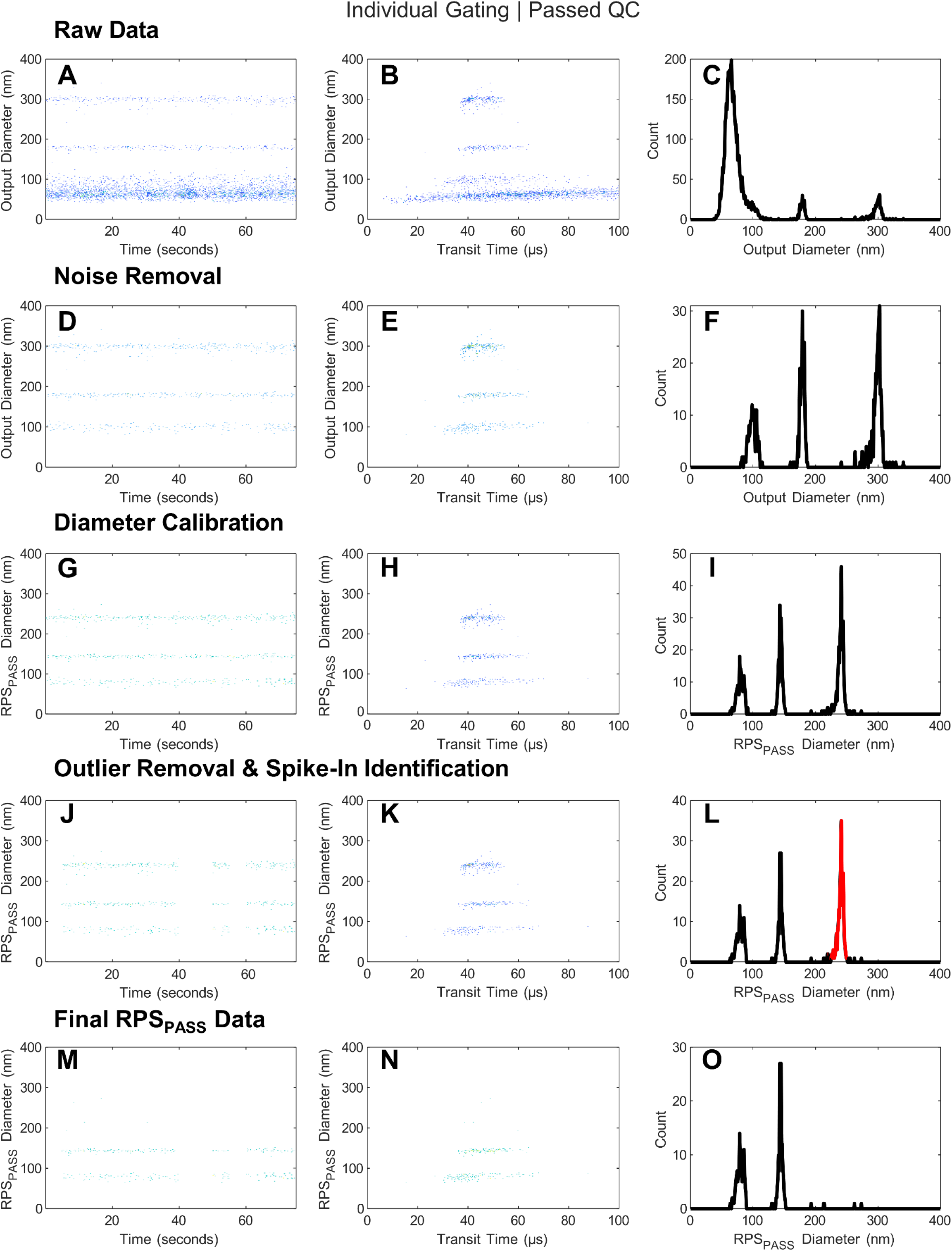
RPS_PASS_ QC plot output. (**A**) Raw nCS1 data plotted using time versus diameter. (**B**) Raw nCS1 data plotted using transit versus diameter. (**C**) Histogram of raw nCS1 data. (**D**) Raw nCS1 data plotted using time versus diameter with noise events removed. (**E**) Raw nCS1 data plotted using transit versus diameter with noise events removed. (**F**) Histogram of raw nCS1 data with noise events removed. (**G**) Dynamically calibrated RPS_PASS_ data plotted using time versus diameter. (**H**) Dynamically calibrated RPS_PASS_ data plotted using transit versus diameter. (**I**) Histogram of dynamically calibrated RPS_PASS_ data. (**J**) Dynamically calibrated RPS_PASS_ data plotted using time versus diameter with outliers removed. (**K**) Dynamically calibrated RPS_PASS_ data plotted using transit versus diameter with outliers removed. (**L**) Histogram of dynamically calibrated RPS_PASS_ data with outliers removed and spike-in beads identified (red). (**M**) Dynamically calibrated RPS_PASS_ data plotted using time versus diameter with outliers and spike-in beads removed. (**N**) Dynamically calibrated RPS_PASS_ data plotted using transit versus diameter with outliers and spike-in beads. (**O**) Histogram of dynamically calibrated RPS_PASS_ data with outliers and spike-in beads removed.

The output of this pipeline starts by taking the raw data (**Fig. 6A-C**) and removing background noise (**Fig. 6D-F**). Upon noise removal, dynamic calibration is performed (**Fig. 6G-I).** Following calibration, outliers are detected and removed using the method shown in **Fig. 5**. The resulting data comprises the particles of interest along with spike-in events (**Fig. 6J-L**). Finally, the spike-in events are detected and gated (**Fig. 6L**), with all events below the spike-in population used to calculate the detectable sample concentration (**Fig. 6M-N**). Gating of the sample population is performed per sample to increase the sensitivity, however, a second global gate that accounts for differing sensitivities across chips is also performed to allow a fair cohort comparison. Output plots for individual sample gates and cohort sample gates are both generated (**Fig. 6O**), with all statistics relating to these gates exported to a spreadsheet.

To investigate the utility of dynamic calibration and cohort gating, a comparison was performed between default nCS1 outputs and RPS_PASS_ outputs using cohort gated human CSF samples from two different groups of donors (**Fig. 7A-B**). Using the cohort gated default output, no significant differences were seen between detected particle counts. However, when RPS_PASS_ dynamic calibration was performed before cohort gating, samples from group 2 were found to have significantly fewer CSF particle counts than those from group 1. No significant differences were found between sample groups when using the default nCS1 output, which showed a larger amount of variation in measurements. While further orthogonal methods are required to understand the differences in particle composition between donor groups, this analysis highlights the utility of increasing the precision and trueness of MRPS measurements for data interpretation and to guide sample comparisons.

**Figure 7:**
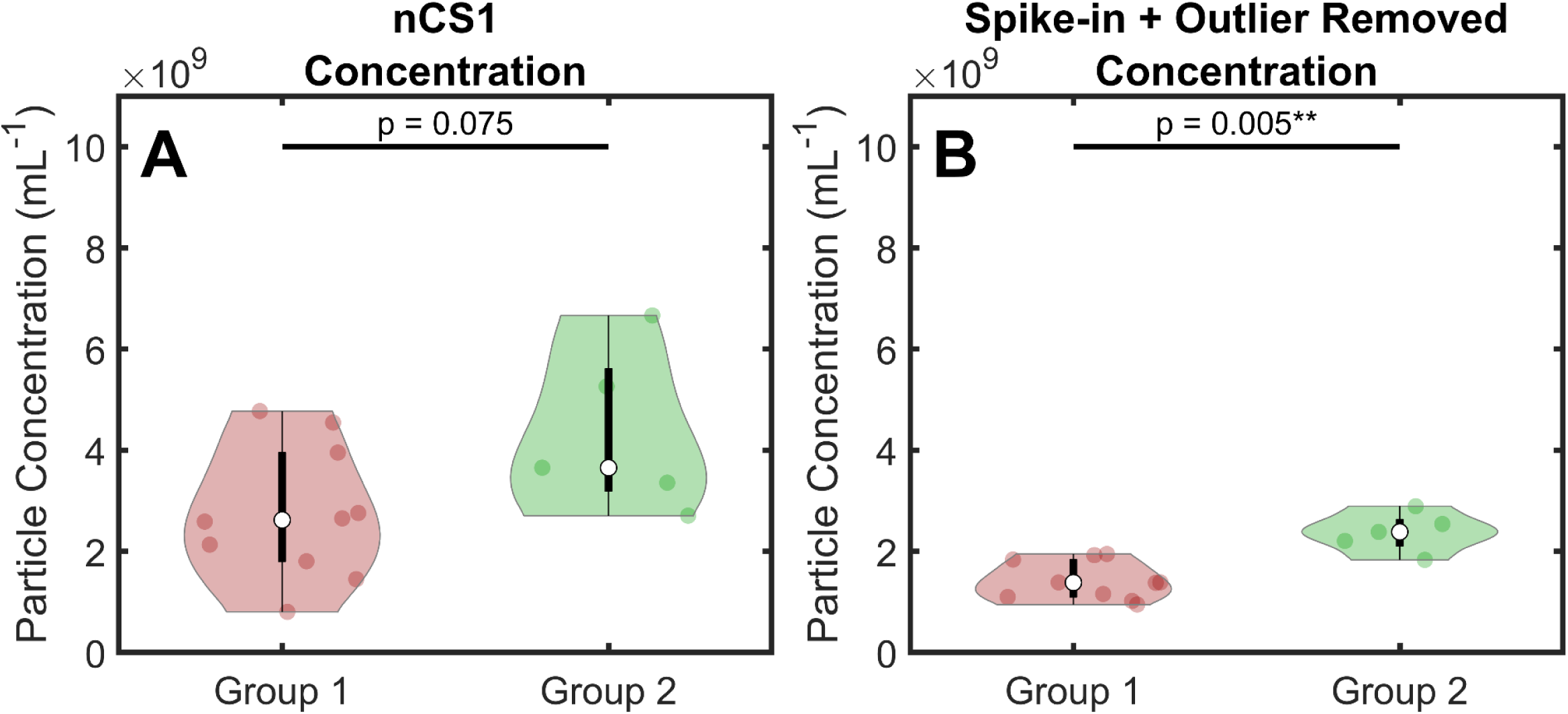
Comparison of default nCS1 data versus RPS_PASS_ data for cohort comparison. (**A**) Cohort gating nCS1 default data comparing particles within CSF from two different groups of donors. (**B**) Cohort gating RPS_PASS_ data comparing particles within CSF from the two groups of donors. Statistical comparison carried out using Wilcoxon signed-rank test.

### Development of a live acquisition interface for improved sample recording

Real-time sample acquisition metrics are essential to ensure reliability of data and to make changes real-time to the sample acquisition to improve the reliability of data for downstream analysis. Information pertaining to the effects of changing parameters such as sample pressure on the transit time of particles through the pore, and visualizing changes in acquisition events, volume, and distributions, are not available. We therefore developed a live acquisition interface for RPS_PASS_ that can process .h5 files as they are written to a folder upon peak-processing by the MRPS software (**Fig. 8**). By using the live acquisition software, it is possible to view RPS_PASS_ calibrated data as soon as it is generated along with metrics such as time, transit time, % CV, acquired volume, and acquired events. Plots of time vs. diameter and diameter vs. transit time are also available immediately upon acquisition. The RPS_PASS_ live acquisition interface allows users to make responsive decisions that allow them to assess and make changes to data during acquisition that will improve the reliability of data in downstream analysis.

**Figure 8:**
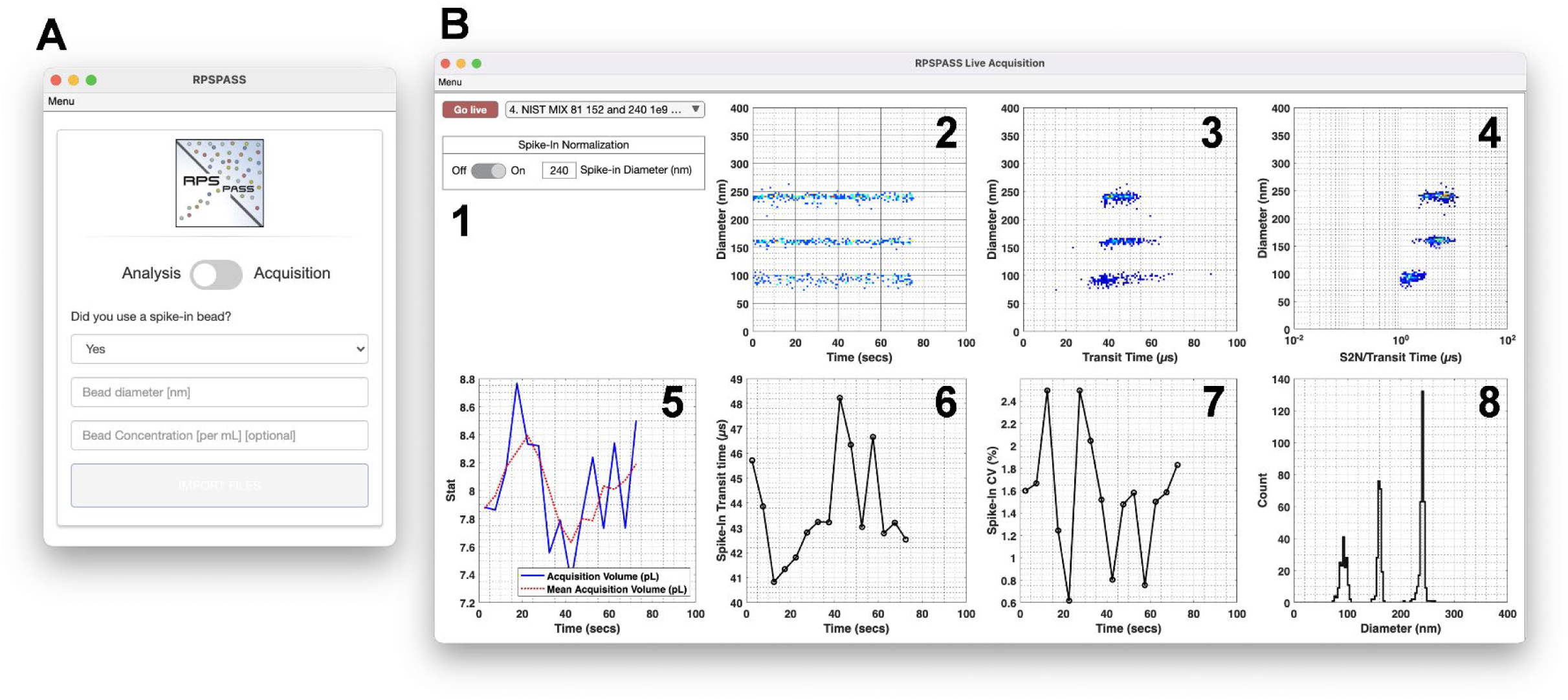
RPS_PASS_ interface. (**A**) Main interface of software allowing for switch between analysis and acquisition modes. **B**) RPS_PASS_ live acquisition interface screenshot showing acquisition of 81, 152, 203 nm NIST-traceable beads. (**B1**) Main user controls to select file to track updates for and live calibration application. (**B2**) Diameter vs. acquisition time plot to track the consistency of data acquisition over time. (**B3**) Diameter vs. transit time plot allows users to monitor resolution consistency. (**B4**) shows the signal-to-noise, transit-time ratio allowing users to gate real particles from noise. (**B5**) shows customizable summary statics plot, allowing users to monitor consistency metrics over time such as acquisition volume and detected events (**B6** & **7**) spike-in transit time and %CV plots over time for monitoring recording consistency and quality. These plots are only shown if a spike-in bead population are used. (**B8**) shows a diameter distribution plot of all events with the option to remove noise events and apply dynamic calibration.

### Developing transparent, standardized MRPS data reporting

Current hurdles to cross-platform data integration are the consistency of reporting detail, data accuracy, and the variety of file types and software available for analyzing data. RPS_PASS_ addresses each of these by performing sample diameter and concentration calibration, performing standardized automated cohort gating, producing a detailed data reporting output, and creating a standard sample metadata table for completion (**Table 2**). To aid in alleviating the lack of software available for MRPS data, files can be normalized and exported in a variety of file formats including .csv, .mat, and .fcs. The conversion of MRPS .hdf5 files to the .fcs file format allows users to access and analyze their data using well-established free and commercial software packages developed for multiparametric flow cytometry data analysis. This is particularly useful given the multiparametric nature of MRPS data which has up to 5 gating parameters, including diameter, signal to noise, symmetry, transit time, and time. The conversion of data to the .fcs file format also aids in the sharing of files to existing public repositories for transparent peer review and joins other freely available software applications that we have developed (i.e. FCM_PASS_^14, 18, 19^ and MPA_PASS_^20^) to facilitate standard EV and nanoparticle analysis, reporting, and data sharing (**Fig. 9**).

**Figure 9:**
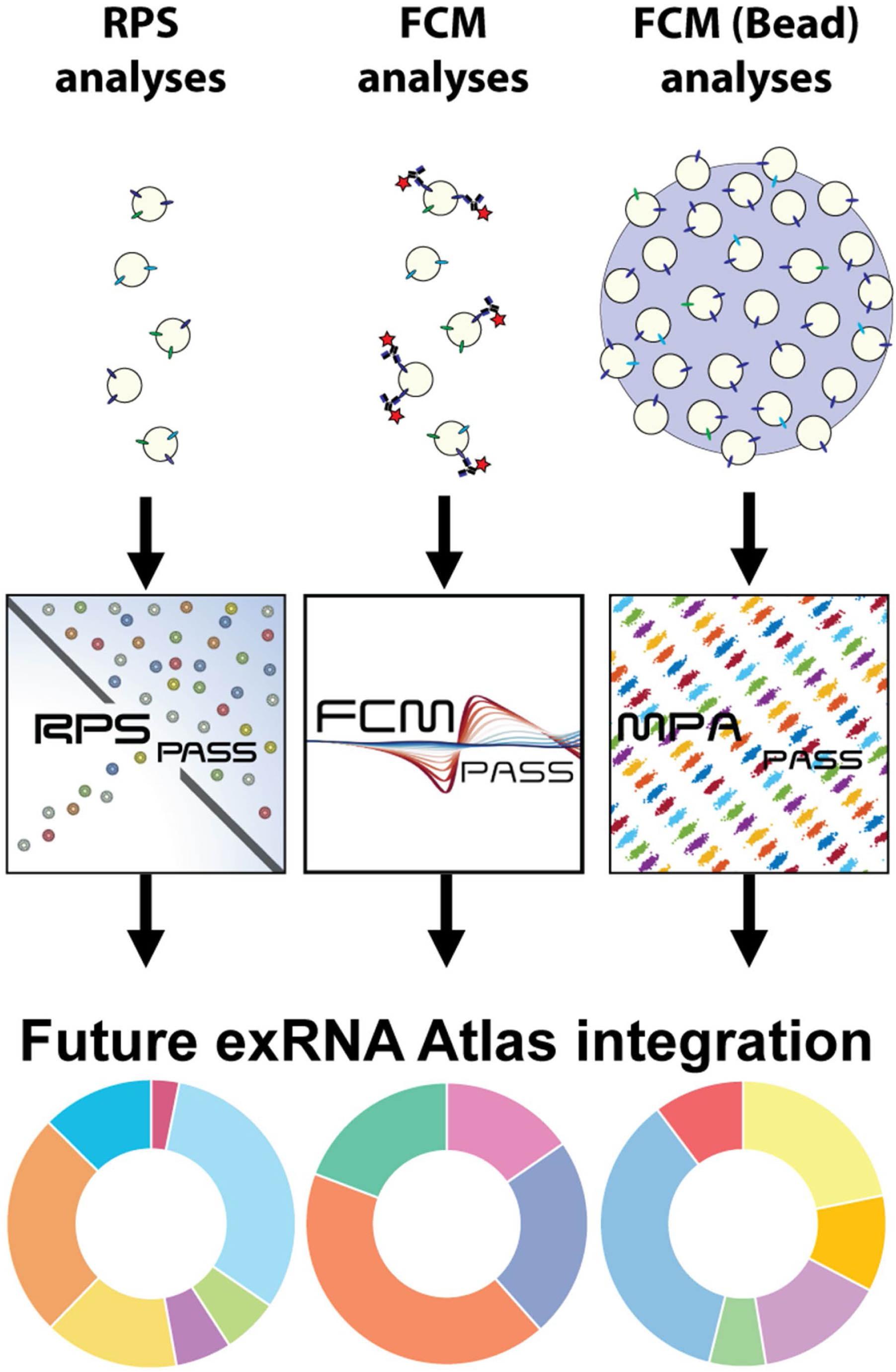
Generation of atlas ready data software packages. A set of software packages have now been developed to enable standardized data analysis and transparent reporting of MRPS data (RPS_PASS_) single particle (FCM_PASS_) and bead-based (MPA_PASS_) flow cytometry data to facilitate repository data integration that can potentially be utilized to develop EV atlas capabilities.

**Table 2.** RPS_PASS_ reporting criteria to improve acquisition and sample transparency.

| Reporting Criteria |
| --- |
| Filename |
| Sample Information |
| Sample Source |
| Sample Isolation |
| Sample Diluent |
| Spike-in Information |
| Cartridge ID |
| Instrument Sheath Fluid |

## CONCLUSIONS

MRPS is a high-throughput platform with high sensitivity and resolution, capable of reporting data calibrated to NIST-traceable standards and supporting reporting of data with quantitative limits of detection. These features have great utility within the EV field as few instruments can detect all EVs, and yet diameter and concentrations are often reported without a limit of detection, which can lead to misinterpretation of results. MRPS is transparent in its .hdf5 file writing, which has allowed for the development of analytical tools such as RPS_PASS_ to increase the power and utility of the platform. With an increase in reporting transparency and an increasing number of techniques capable of generating single EV data in calibrated units, it is now possible to start integrating cross-platform standard unit data. The aggregation of this type of data in the literature will inevitably aid in further understanding of EVs and other extracellular nanoparticles and make it possible for centralized repositories or atlases to form where inter-laboratory data generation can be aggregated and cross-analyzed. We have previously demonstrated the utility of MRPS orthogonal measurements with high-sensitivity flow cytometry to derive single-vesicle refractive index derivations.^21^

Like light-scatter based NTA measurements, the specificity of MRPS particle measurements is dependent upon the purity of the sample for the particles of interest. The use of MRPS in combination with other orthogonal methods that can provide greater specificity, such as fluorescence-based single particle analysis methods or high-resolution imaging methods, is therefore recommended. RPS_PASS_ has also been able to aid with the derivation of effective refractive index by utilizing the fluorescent properties of rEVs, which are proportional to their surface area, and by calibrating their light scattering signal to standard units of scatter cross section.^21^ In this work we utilized CSF EVs and rEVs buffered with 1% Tween 20 for our MRPS measurements. **Supplemental Figure 2** illustrates that at this concentration of Tween we successfully buffer rEVs without lysing of vesicles, however it should be mentioned that this may not hold true for all types of EVs, some subsets of which may be more sensitive to detergent lysis.

Use of RPS_PASS_ increases the accuracy of data outputs from MRPS instruments and aids in robust cohort analysis by removing outliers, noise, and drawing gates that are consistent across samples while maximizing sensitivity. Along with automated analysis, the use of RPS_PASS_ provides dexterity for users to explore, analyze, and share their data in different formats. RPS_PASS_ also focuses on increasing measurement and analysis transparency by outputting summary QC plots for all samples and aggregating all sample statistics and gating criteria in a single spreadsheet. In similar fashion to MIFlowCyt-EV for flow cytometry data,^15^ we have proposed a set of key meta-data criteria to be reported for each sample acquisition in the RPS_PASS_ output spreadsheet. In doing so, we believe that researchers will be more likely to comply and therefore the transparency of MRPS data in the field will improve, facilitating others to perform validation experiments.

## Author Contributions

MLP, JCJ, and JAW contributed towards the conceptualization, data curation, methodology, formal analysis, and manuscript writing, review, and editing. MLP, SC, BK, DAJ, TT, EHS, MN, and JAW contributed to investigation, resources, and validation. JAW contributed to the software development, and validation of data. JCJ, JAW, and AM contributed towards project administration and supervision. All authors contributed to manuscript writing, revision, and project organization and coordination. JCJ, IG, AH, SJ and AM contributed to funding acquisition. JCJ, AH, CP, and SJ contributed towards resources. JEA, MR, and AM contributed to data curation.

## Conflicts of interest

MLP, SC, BK, TT, DAJ, EHS, VF, CP, JEA, JS, MR, AM, IG, and SJ declare no conflicts of interest. JAW and JCJ are inventors on NIH patents and patent applications related to EVs. AH is an inventor on a UGent patent on rEV technology (WO2019091964).

## Acknowledgements

The authors would like to thank Spectradyne for their help providing the authors with a detailed understanding of the nCS1 platform at a hardware and software level. JAW, SC, DJ, JS, and JCJ were supported by the Intramural Research Program of the National Institutes of Health (NIH), National Cancer Institute, and Center for Cancer Research. MLP was a National MS Society (NMSS) postdoctoral fellow (grant FG-2107-38321). JCJ acknowledges NIH ZIA BC011502, NIH ZIA BC011503, and a Prostate Cancer Foundation Young Investigator Award. JAW was an International Society for Advancement of Cytometry (ISAC) Marylou Ingram Scholar 2019-2023. JCJ, IG, MR and AM acknowledge NIH 4UH3TR002881. MR and AM acknowledge NIH 5U54DA049098.

## Supplemental Figures

**Supplemental Figure 1:**
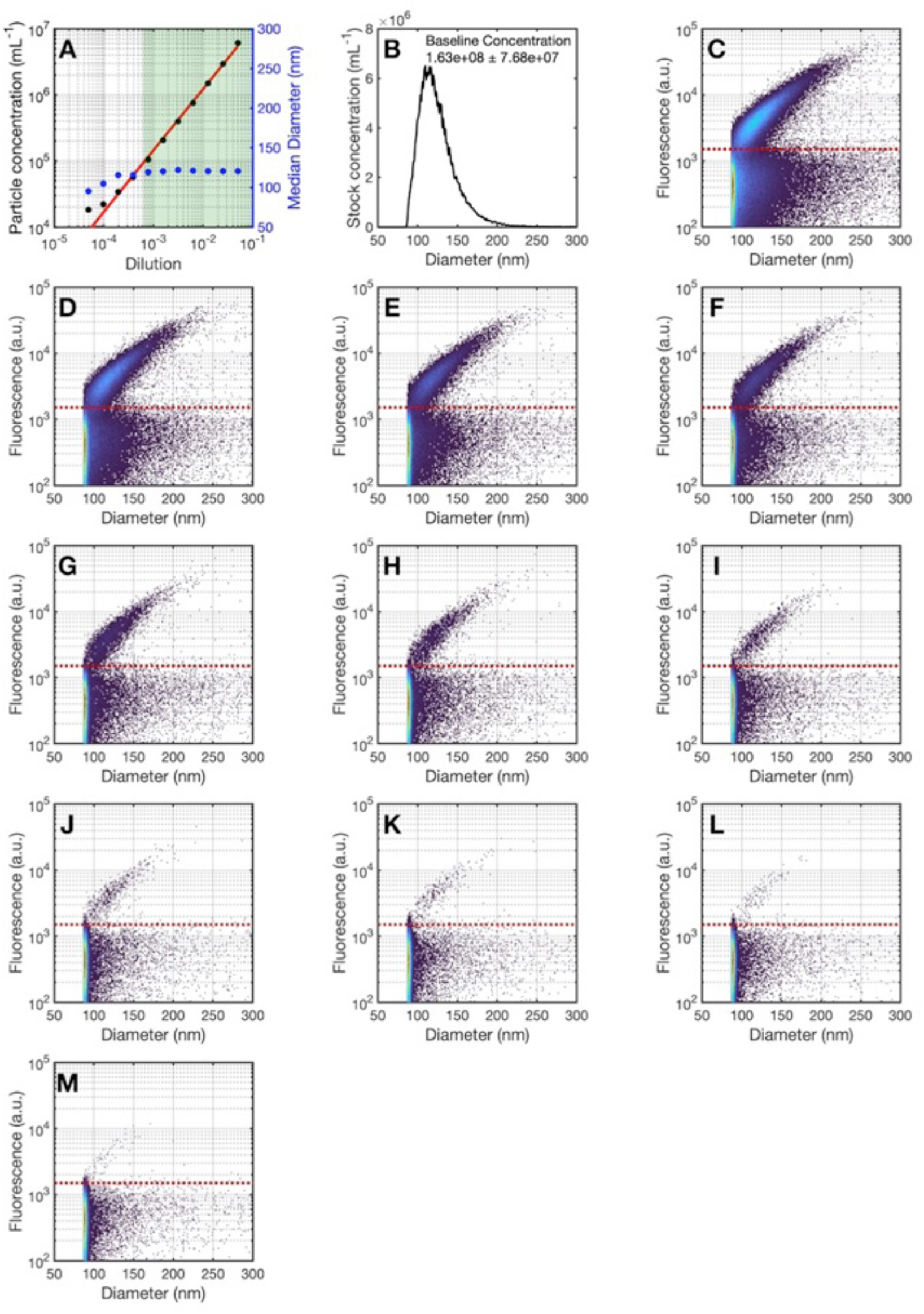
Flow cytometry quality control data for rEV acquisition. **A**) serial dilution of rEVs with concentration vs. dilution factor (black dots, red regression line) and median light scatter derived diameter (blue dots). Green shaded area shows the most reliable concentration dynamic range of detection for rEVs. **B**) diameter distribution of rEVs using light scatter calibration with the detected concentration. **C-M**) show raw data plots from low dilution factor to high dilution factor.

**Supplemental Figure 2:**
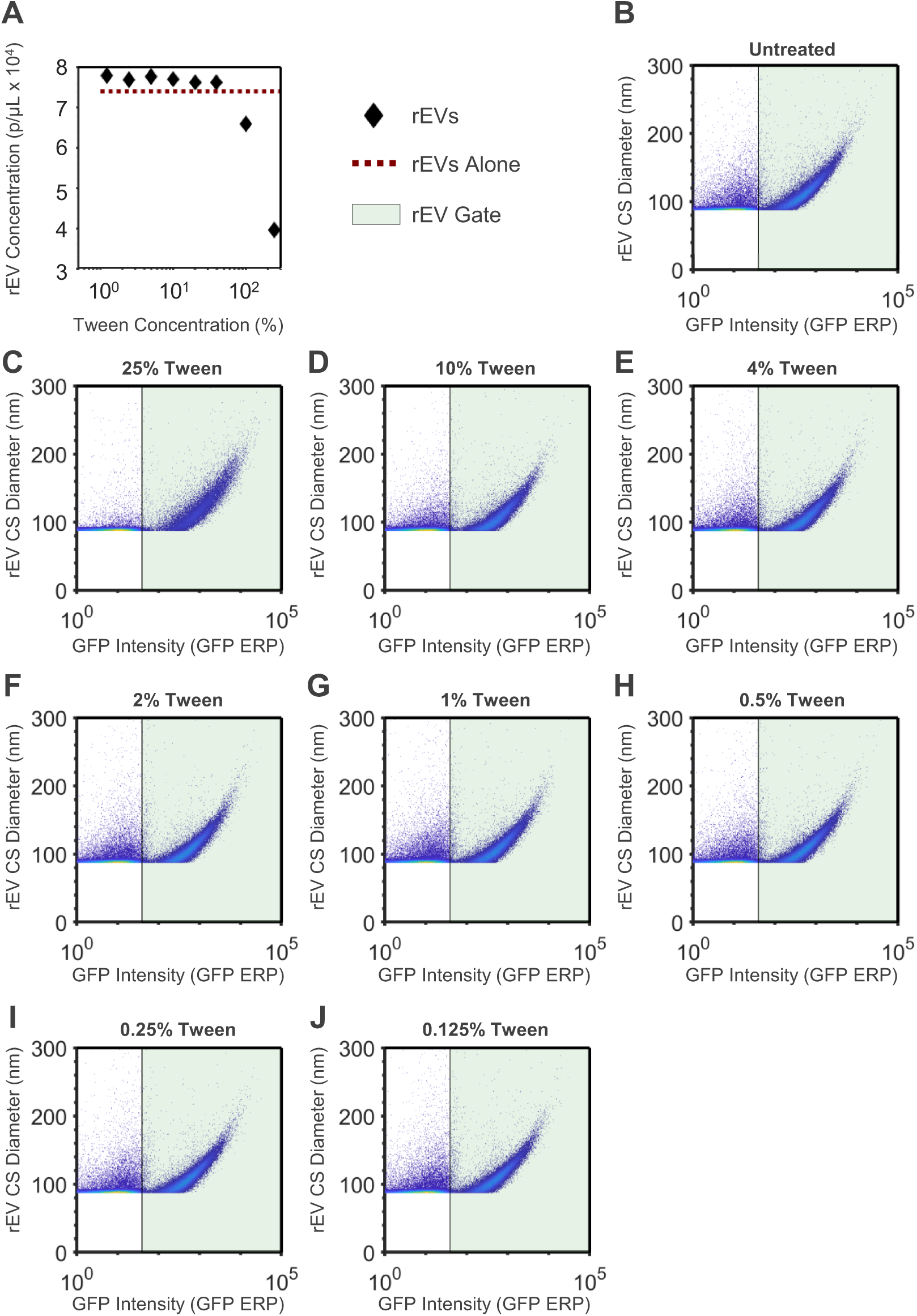
Tween titration control data for rEV acquisition. **A**) Summary flow cytometry data of rEVs (black dots) with an increasing titration of Tween 20. rEVs alone with no Tween 20 treatment are shown (red dotted line). **B-J**) Raw data plots of each rEV +/- Tween 20 treatment flow cytometry acquisitions. Green shaded areas show the selected rEV gate.

## Notes

### Competing Interest Statement

The authors have declared no competing interest.

